# Experimental evolution of *Plasmodium yoelii* in single and helminth-coinfected mice

**DOI:** 10.1101/2025.10.22.683851

**Authors:** Aloïs Dusuel, Luc Bourbon, Emma Groetz, Mickaël Rialland, Benjamin Roche, Bruno Faivre, Gabriele Sorci

**Author notes:** These authors contributed equally to this work (Aloïs Dusuel, Luc Bourbon).

## Abstract

Coinfection has the potential to affect key traits describing the infection dynamics, the severity of the disease and *in fine* parasite fitness. However, despite its pervasiveness, experimental work investigating how parasites adapt to the conditions provided by a coinfected host is mostly missing. Here, we adopted an experimental evolution approach to investigate if coinfection with the nematode *Heligmosomoides polygyrus* (Hp) affected the infection dynamics and virulence of the murine malaria parasite *Plasmodium yoelii* (Py). To this purpose, lines of Py were passaged either in single infected hosts (SI-lines) or in hosts that had been previously infected with Hp (COI-lines). After five and seven passages, the infection dynamics and virulence of evolved lines were compared to the ancestral Py population during single infection trials. As expected, we found that serial passages increased parasitemia and Py virulence, due to the competitive advantage of genotypes with the fastest replication rate, but SI-lines and COI-lines had similar replication rate and virulence. Hosts infected with evolved lines of Py were also less tolerant (steeper slope between red blood cell counts and parasitemia) but again there was no difference between SI-lines and COI-lines. In a second experiment, COI-lines were also used to infect hosts during coinfection trials, allowing us to compare within-host Py replication when the environment during the evaluation trials matched the environment experienced during the passages and when they were mismatched. The results showed that COI-lines used during single infection trials (mismatched environment) had a slower replication rate compared to both SI-lines and COI-lines used during matched-environment trials. Overall, although we did not find any difference in the virulence of SI-lines and COI-lines after seven passages, Py rapidly adapted to the environmental conditions provided by single infected or coinfected hosts, as shown by the slower replication rate found in mismatched-environment trials.

**Author Summary:** Coinfection can alter infection dynamics, disease severity, and parasite fitness, yet experimental evidence on parasite adaptation to coinfected hosts is scarce. Here we used experimental evolution to assess whether coinfection with the nematode *Heligmosomoides polygyrus* influences the infection dynamics and virulence of the murine malaria parasite *Plasmodium yoelii*. Parasite lines were serially passaged in either single infected hosts or hosts previously infected with *Heligmosomoides polygyrus*. After multiple passages, evolved lines exhibited increased parasitemia and virulence compared to the ancestral strain, but no significant differences were found between lines evolved in single or coinfected hosts. Notably, parasite replication was reduced when lines evolved in coinfected hosts were tested in single infection environments, indicating rapid adaptation to host infection context. This suggests that while coinfection shapes within-host conditions, *Plasmodium* adapts rapidly to these conditions. These results have possible implications for understanding parasite evolution in fluctuating host environments, and the microevolutionary consequences for malaria of the deworming campaigns that are implemented to control helminthiases.

## Introduction

Why parasites kill their hosts has puzzled biologists for decades (Zampieri et al. 2025). Indeed, parasites (general term including both micro and macroparasites) usually rely on host survival to produce propagules, and host death inevitably incurs the end of the infectious period (the period during which parasites produce transmissible stages). Based on this simple idea, it was believed that parasite virulence (the degree of infection-induced damage potentially leading to host death) was a maladaptive trait and that selection should consistently favor more benign parasite strains (Smith’s law of declining virulence, Méthot 2012). This view was nevertheless at odds with the observation that some parasites have maintained high levels of virulence over time. A solution to the puzzle was finally provided by the seminal work of Anderson and May (1979, 1981, 1982), who suggested that parasite virulence can evolve up and down depending on the balance between the benefits arising from increased host exploitation (i.e., increased within-host replication rate) and the costs arising from host death (i.e., decreased infectious period). Since then, this trade-off model of virulence evolution has attracted considerable attention both from theoreticians and empiricists who have investigated the assumptions of the model and the generality of the predictions (e.g., Frank 1996; Wickham 2007; Cressler et al. 2016; Clay and Rudolf 2019).

Whatever the direction of the selection on virulence, parasites are expected to adapt to the prevailing environmental conditions, as to maximize transmission success (e.g., Mackinnon and Marsh 2010, Rono et al. 2018). Experimental evolution in the lab has consistently shown that parasites passaged in a novel host become adapted to it while becoming maladapted to the ancestral host (Ebert 1998, Rafaluk et al. 2015). Evidence of parasite adaptation to novel hosts comes also from natural outbreaks. For instance, during the Ebola virus outbreak in west Africa, the glycoproteins allowing the viral entrance in the host cells increased their affinity towards the human receptor while losing affinity towards the ancestral bat receptor (Urbanowicz et al. 2016), indicating ongoing viral adaptation to the human host.

In natural populations, hosts are exposed to a multitude of parasites that can concomitantly infect the same individuals (Viney and Graham 2013). Coinfection has the potential to deeply change the environment encountered by parasites compared to single infected hosts for several reasons (Graham 2008, Jolles et al. 2008, Ezenwa and Jolles 2011, Shen et al. 2019, Desai et al. 2021, Petrellis et al. 2023, Godinho et al. 2024, Fogang et al. 2025). Indeed, depending on the nature of the coinfecting parasites, the adaptive response of the host can be polarized towards different immune effectors (Th1/Th17 vs Th2), making the ground more or less favorable for host exploitation (Supali et al. 2010, Ma et al. 2020). In addition to this, parasites (e.g., helminths) can also directly interfere with the host immune response and the associated immune modulation can promote the infection by competing parasites (Hartgers and Yazdanbakhsh 2006, Salgame et al. 2013). Finally, coinfecting parasites might directly compete for resources provided by the host, through bottom-up regulatory mechanisms (Budischak et al. 2018).

Malaria is still one of the deadliest infectious diseases worldwide, with an estimated number of about 600,000 deaths in 2023 (WHO 2025). Malaria endemicity largely overlaps with the zones where helminthiases (soil-transmitted helminth infections) are also highly prevalent. As a consequence, coinfection between malaria and helminths is common among the poorest populations living in the intertropical area (Kepha et al. 2015, Afolabi et al. 2021). Several epidemiological studies have investigated the effect of the coinfection on the severity of malaria symptoms with somehow mixed results since both aggravating and protective effects have been reported (Degarege and Erko 2016, Hürlimann et al. 2019, Masoud et al. 2024). Although soil-transmitted helminths do not exert the same mortality burden as malaria, they represent a public health concern due to the high prevalence of infection and their effect on infant physical and cognitive development. Massive deworming is one of the control strategies that is regularly implemented to reduce the burden of infection with soil-transmitted helminths (Welch et al. 2017), but its efficacy and rationale has been debated (Humphries et al. 2012, Taylor-Robinson et al. 2019), on the ground, among others, of the risk of emergence of drug resistant strains (Tinkler 2020, Coffeng et al. 2024). Moreover, another aspect linked to deworming that has been barely considered is the possible feedback on coinfecting microparasites (e.g., malaria) and their adaptive response to the clearance of helminth infection. In one of these few attempts, Fenton (2013) built a model where he analyzed the possible outcomes of deworming on two key traits of a hypothetical coinfecting microparasite, the basic reproduction number and virulence. The model explicitly considered different types of within-host interactions between worms and pathogens, where for instance worms aggravate the severity of the infection with the microparasite, or improve host recovery rate. The findings suggested that the outcome of deworming is largely context-dependent, according to the type of within-host interaction. In particular, when helminths aggravate the severity of the pathogen infection (increase in host mortality), deworming should select for increased pathogen virulence. Similarly, when helminths weaken the host recovery rate, the model predicted that deworming should promote pathogen virulence, at least when the natural worm burdens are low to moderate.

Here, we wished to investigate if malaria parasites can adapt to the conditions provided by a coinfected host using a model involving two murine parasites, *Plasmodium yoelii* (hereafter Py) and the gastrointestinal nematode *Heligmosomoides polygyrus* (hereafter Hp). In a recent paper, we showed that Py incurs higher costs when infecting hosts already infected with Hp (compared to the single infection) (Dusuel et al. 2025). We also showed that the changes in the host immune response induced by Hp promoted Py replication and slowed down host recovery rate (Dusuel et al. 2025). Therefore, when mosquitoes bite and transmit Py to a host already harboring a Hp infection, Py will likely experience a very distinct immune environment compared to the one provided by a non-infected host. Here, we adopted an experimental evolution approach where Py was passaged either in single infected (potentially reproducing the environment provided by a dewormed host) or in coinfected (with Hp infecting first) hosts, in a one-sided evolution design. After five and seven passages, the evolved Py lines were used to infect mice in single infection trials and the symptoms elicited by the infection were compared to those elicited by the ancestral Py population. We measured several proxies of disease severity, including traits related to the host capacity to tolerate the cost of the infection. In addition to this, we also ran a second experiment where Py lines evolved in coinfected hosts were also used in coinfection trials. This allowed us to investigate whether lines passaged and tested in the same environmental conditions (matched environments) fared better than lines experiencing mismatched environments. This experiment therefore attempted to reproduce the conditions encountered by *Plasmodium* potentially switching from wormed to dewormed hosts.

## Materials and Methods

### Ethical statement

Experiments were approved by the Comité d’éthique Grand Campus Dijon of the University of Burgundy and were authorized by the French Ministry of Higher Education and Research under the numbers 33492 and 40849.

### Experimental animals

C57BL/6JRj female mice (7-10 weeks old) were purchased from Janvier Labs (Le Genest-Saint-Isle, France), housed in cages containing 5 individuals under pathogen-free conditions, and maintained under a constant temperature of 24°C and a photoperiod of 12h:12h light:dark with *ad libitum* access to water and standard chow diet (A03-10, Safe, Augny, France). All mice were acclimatized to the housing conditions during, at least, 7 days prior to the start of the experiments, were monitored twice a day to check health status, and euthanized by cervical dislocation under anesthesia with isoflurane either if they reached previously defined end points (and considered as mortality events in the survival analysis) or at specific days post-infection (p.i.) for terminal collection of blood and spleen (and considered as censored events in the survival analysis).

### Serial passages

At the beginning of the experiment, one group of mice was infected with *Plasmodium yoelii* 17XNL by intra-peritoneal (i.p.) injection with 5x10^5^ infected red blood cells (iRBC) suspended in 0.1 ml of PBS (single infection group), and another group was first infected with *Heligmosomoides polygyrus bakeri* by oral gavage with L3 larvae (300 larvae suspended in 0.2 ml of drinking water) and after 28 days infected with Py as described above (coinfection group). Infected red blood cells used for single infections and coinfections came from the same stock.

At day 14 post Py infection, Py from the single infection group was passaged to five mice that received it as a single infection (5x10^5^ iRBC) and Py from the coinfection group was passaged to five mice that had been previously infected with Hp (28 days prior to the Py infection). Therefore, we had two treatments: either Py was passaged in single infected mice (SI-lines) either in coinfected mice (COI- lines), each treatment having 5 replicated lines (5 mice). Infection with Hp was done using the stock population of nematodes, in other words, only Py was allowed to respond to the treatment over the passages, in a one-sided evolution design. The setup was repeated for 10 passages. After 5, 7 and 10 passages, the evolved lines of Py were used to infect 15 naïve mice (single infection trials) whose health status and parasitemia was monitored during 14 days post Py infection. This experiment allowed us to compare the infection dynamics and virulence of evolved lines compared to the ancestral one depending on the selection regime (the passage treatment): single infection vs. coinfection. Due to the mortality during the passages, the experiment was stopped after 7 passages for the COI-lines.

In a second experiment, the COI-lines were also used for coinfection trials (mice that had been previously infected with Hp at day -28). Therefore, we could compare the performance of Py when encountering the same environmental conditions as those experienced during the passages (SI/si or COI/coi) to the performance of Py encountering mismatched environments (COI/si) (capital letters indicating the environment during the passages and the small letters indicating the environment during the trials). Figure 1 schematically illustrates the experimental design.

**Figure 1.**
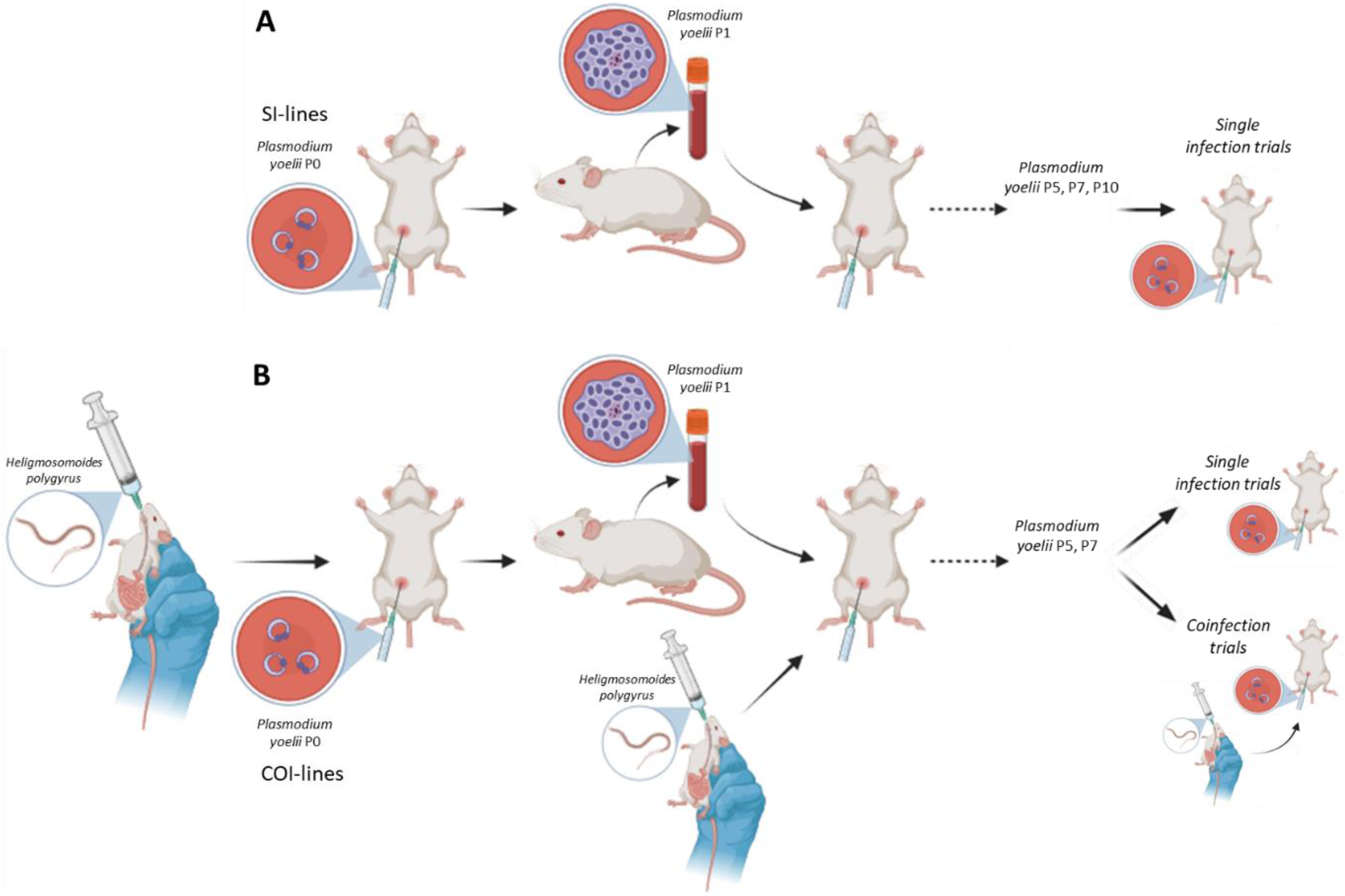
Schematic view of the experimental evolution setup. Py was serially passaged in single infected (A, SI-lines) or coinfected hosts (B, COI-lines). In the coinfected treatment, hosts were infected with Hp 28 days prior to the infection with Py. Five mice were infected per passage, providing five replicated transmission lines. Py was passaged at day 14 post-infection. Only Py was passaged, Hp coming from the stock population (i.e., only Py was allowed to evolve). After 5, 7 and 10 passages, SI-lines were tested during single infection trials. After 5 and 7 passages, COI-lines were tested during both single infection and coinfection trials. Please note that although the figure depicts albino mice, the experiment involved C57BL/6J mice. Figure created in BioRender.

### Monitoring of health status, hematological and plasma parameters, and Py parasitemia

Mice were weighed at day 0, 3, 7, 9 and 14 post Py infection (± 0.1 g) and body mass was used as an integrative marker of host health. Red blood cells (RBCs) were counted at day 0, 3, 9, 14 post Py infection using a SCIL Vet abc Plus+ hematology analyzer (Horiba Medical, Montpellier, France) on 10 µl of blood collected from the tail tip and disposed in 0.5 ml tubes with 1 µl of heparin (Héparine Choay® 25 000 UI/5ml, Sanofi-Aventis, Gentilly, France). Parasitemia was assessed at day 3, 7, 9, and 14 post Py infection by smearing a drop of blood on a slide. Slides were fixed with methanol and then stained with a Giemsa RS solution (Carlo Erba Reagents, Val de Reuil, France) (10% v/v in phosphate buffer) for 15 minutes and rinsed with water. Infected RBCs were counted using an optical microscope (Eclipse E600, Nikon Corporation, Tokyo, Japan) under magnification x1000, and parasitemia expressed as the proportion of iRBCs.

At day 3 and 14 post Py infection, five and ten mice per treatment, respectively, were euthanized by cervical dislocation under isoflurane anesthesia, for terminal collection of total peripheral blood and spleen. Plasma was separated from total blood by centrifugation (7 min 3000 rpm 4°C), aliquoted and stored at -80°C. Spleen was removed, about one third was cut and immediately frozen in liquid nitrogen for RT-qPCR, and the remaining was kept on ice in PBS before the flow cytometry staining procedure.

Plasma levels of mouse erythropoietin (EPO) and heme oxygenase-1 (HO-1) were assessed with EM28RB ELISA kit from Invitrogen™ (Carlsbad, California, USA) and ab229431 ELISA kit from Abcam (Cambridge, UK), respectively. Plates were read with a SpectraMax iD3 multi-mode microplate reader (Molecular Devices, San Jose, California, USA).

### RNA extraction and qPCR for IFN-γ, IL-10 and TGF-β1 gene expression

Spleen tissue was homogenized in TRIzol™ Reagent (Invitrogen™, Carlsbad, California, USA) under strong agitation using 0.5 mm glass beads and a Precellys® 24 Touch homogenizer (Bertin Technologies, Montigny-le-Bretonneux, France). RNA extraction was performed following the manufacturer’s instructions. RNA concentration was measured with a NanoPhotometer® N50 (Implen, Munich, Germany). Reverse transcription was performed with a High-Capacity cDNA Reverse Transcription Kit (Applied Biosystems™, Foster City, California, USA) from 1.5 µg total RNA. Quantitative PCR was performed with PowerUp™ SYBR™ Green Master Mix (Applied Biosystems™) on a QuantStudio™ 3 Real-Time PCR System (Applied Biosystems™). We used two housekeeping genes (β-actin and GAPDH); β-actin provided more consistent values (less intra-group variability) and therefore we used it as the reference gene. Primer sequences are reported in the supplementary material (table S1), and were checked for specificity using the Primer-BLAST tool (NCBI). Melt curve analysis was also performed in all samples to check for the presence of nonspecific amplification products.

### Assessment of FoxP3^+^ Treg cells

Spleens were homogenized with a 70 µm cell strainer and washed two times with PBS to obtain a single-cell suspension. Red blood cells were removed from suspensions with eBioscience™ RBC Lysis Buffer (Invitrogen™) (3 min incubation at room temperature) and washed two times with PBS before staining procedure.

Live cells were counted with viability trypan blue dye and dispatched in 96-well V bottom plates to obtain 1.10^7^ cells/well. Cells were then stained with LIVE/DEAD™ Fixable Blue Dead Cell Stain Kit (Invitrogen™) in PBS for 30 min at 4°C, incubated in stain buffer with BD Fc Block™ (BD Pharmingen™, Franklin Lakes, New Jersey, USA) for 15 min at RT (0.5 µg/well), and stained with surface antibodies in brilliant stain buffer (see table S2 for details on the markers used) for 30 min at 4°C. Cells were then fixed with eBioscience™ Foxp3 / Transcription Factor Staining Buffer Set (Invitrogen™) for 40 min at RT prior permeabilization and intracellular staining in brilliant stain buffer (table S2) for 30 min at RT. Cells were resuspended in stain buffer, filtered with 100 µm nylon mesh and analyzed on a 4-laser Cytek Aurora™ spectral flow cytometer (Cytek® Biosciences, Fremont, California, USA) in the ImaFlow facility part of the US58 BioSanD (Dijon, France). Events were gated and analyzed with SpectroFlo® v3.0.0 software (Cytek® Biosciences) (see figure S1 for the gating strategy).

### Statistical analyses

For the first experiment, for each phenotypic trait, we computed the changes in evolved lines relative to the ancestral population for the two selection regimes (SI-lines and COI-lines) as the ratio between the individual value after passage 5 or 7 and the mean of the ancestral population. Therefore, values of 1 indicate no phenotypic changes in the evolved lines compared to the ancestral population. For the IFN-γ and IL-10 gene expression, we computed the ΔΔ Ct as the matched day of sampling difference between the Δ Ct of the individual values of SI-lines and COI-lines and the mean Δ Ct of the ancestral population (0 indicating no phenotypic changes in the evolved lines compared to the ancestral population). These values were analyzed using a two-step procedure. First, we ran intercept only general linear mixed models (GLMM) to test if values of the evolved lines were different from the ancestral population (intercept different from 1, or 0 for ΔΔ Cts) at day 14 post-infection. These models included the transmission line identity as a random factor. Second, we ran GLMMs to test if values were different between evolved lines. These models included mouse identity (for traits with repeated measurements per individual) and transmission line identity as random effects, the selection regime (SI-lines, COI-lines), the passage (5 or 7), and time post Py infection (for traits that were measured at different time points), and the two-way and three-way interactions as fixed effects. We systematically checked whether models including a squared time p.i. term better fitted the data, and only left it in the model if it was the case (i.e., models on parasitemia).

Differences in survival between groups were analyzed using the Kaplan Meier estimator and significance assessed using a Log-Rank test. Adjustment for multiple comparisons for the Log-Rank test was done using Sidak adjusted p values.

For experiment 2, we directly compared the parasitemia of SI-lines tested in si-hosts, COI-lines tested in coi-hosts (the matched environment groups) and COI-lines tested in si-hosts (the mismatched environment group), using a generalized linear mixed model with a beta distribution of errors (the dependent variable being the proportion of iRBC). The model included mouse identity and the transmission line identity as random factors and time post Py infection, squared time post Py infection, trial group (SI/si, COI/si, COI/coi), plus the two-and three-way interactions as fixed effects. We also ran a second model where the two matched-environment groups (SI/si and COI/coi) were clustered together and compared to the mismatched-environment group.

Differences in sample size across groups reflect missing values, either due to the reaching of the end point or to technical issues during the processing of the sample.

All the statistical analyses and figures were done using SAS Studio.

## Results

### 1) *Plasmodium* evolution in single infected and coinfected hosts

#### a) Disease severity in evolved Py lines

Body mass loss is an integrative proxy of disease severity during malaria infection. We, therefore, first compared how changes in body mass, relative to mice infected with the ancestral Py population, varied over time for mice infected with SI-lines or COI-lines. The GLMM reported a highly significant negative effect of time p.i., with values crossing the line of no difference with respect to the ancestral Py population, indicating that over the course of the infection mice infected with the evolved Py lines lost more body mass (table 1, figure 2a). An intercept only GLMM that included data at day 14 p.i. showed that values were significantly lower than 1, as shown by the 95% confidence intervals non overlapping 1 [intercept (95% CI) = 0.907 (0.846/0.968), n = 48]. The model also indicated that the slope relating changes in body mass and time p.i. was steeper for the SI-lines and for lines tested after 5 passages (table 1), but parameter estimates and visual inspection of figure 2a,b suggested very modest effect sizes.

**Figure 2.**
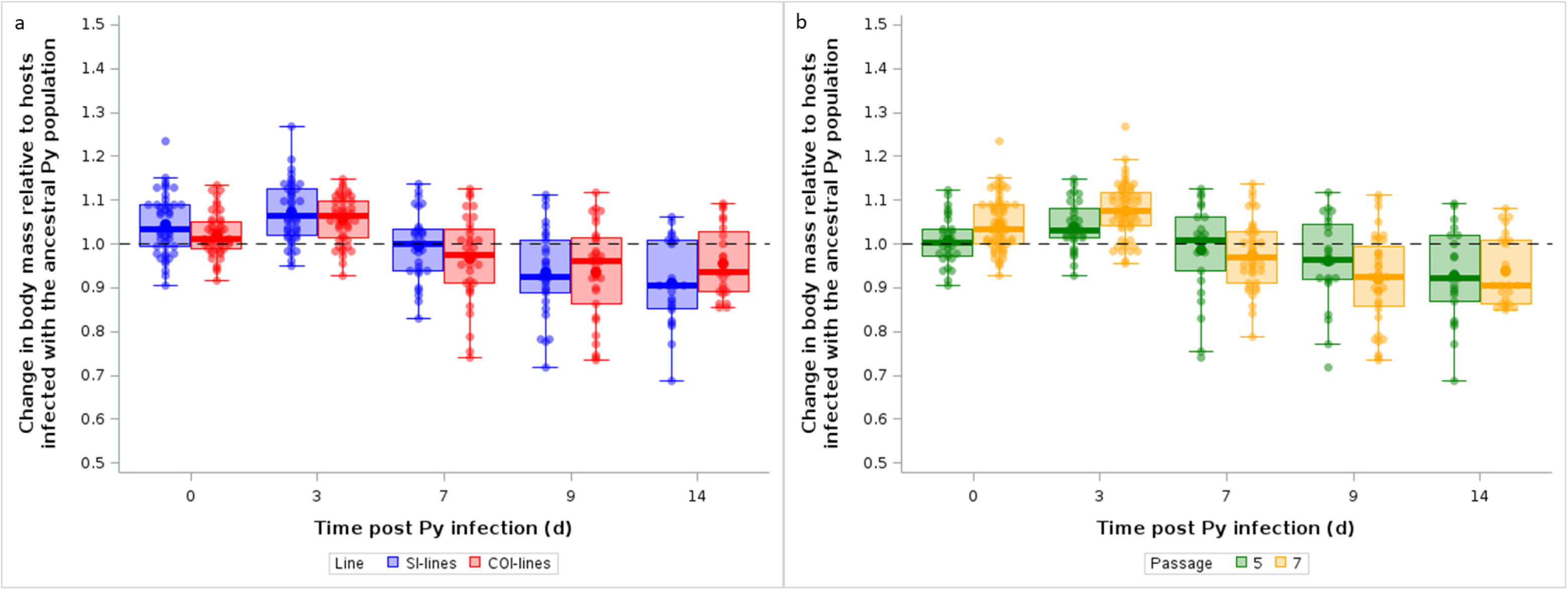
Changes in body mass over the course of the infection during single infection trials. Body mass is expressed as change relative to hosts infected with the ancestral Py population; a) Changes in body mass for hosts infected with SI-lines vs. COI-lines; b) Changes in body mass for hosts infected with Py after 5 vs. 7 passages. The dotted line represents no change with respect to hosts infected with the ancestral Py population. Dots represent the raw data, the boxes represent the interquartile range (IQR), the horizontal lines the median, and whiskers the range of data within 1.5 the IQR.

**Table 1.**
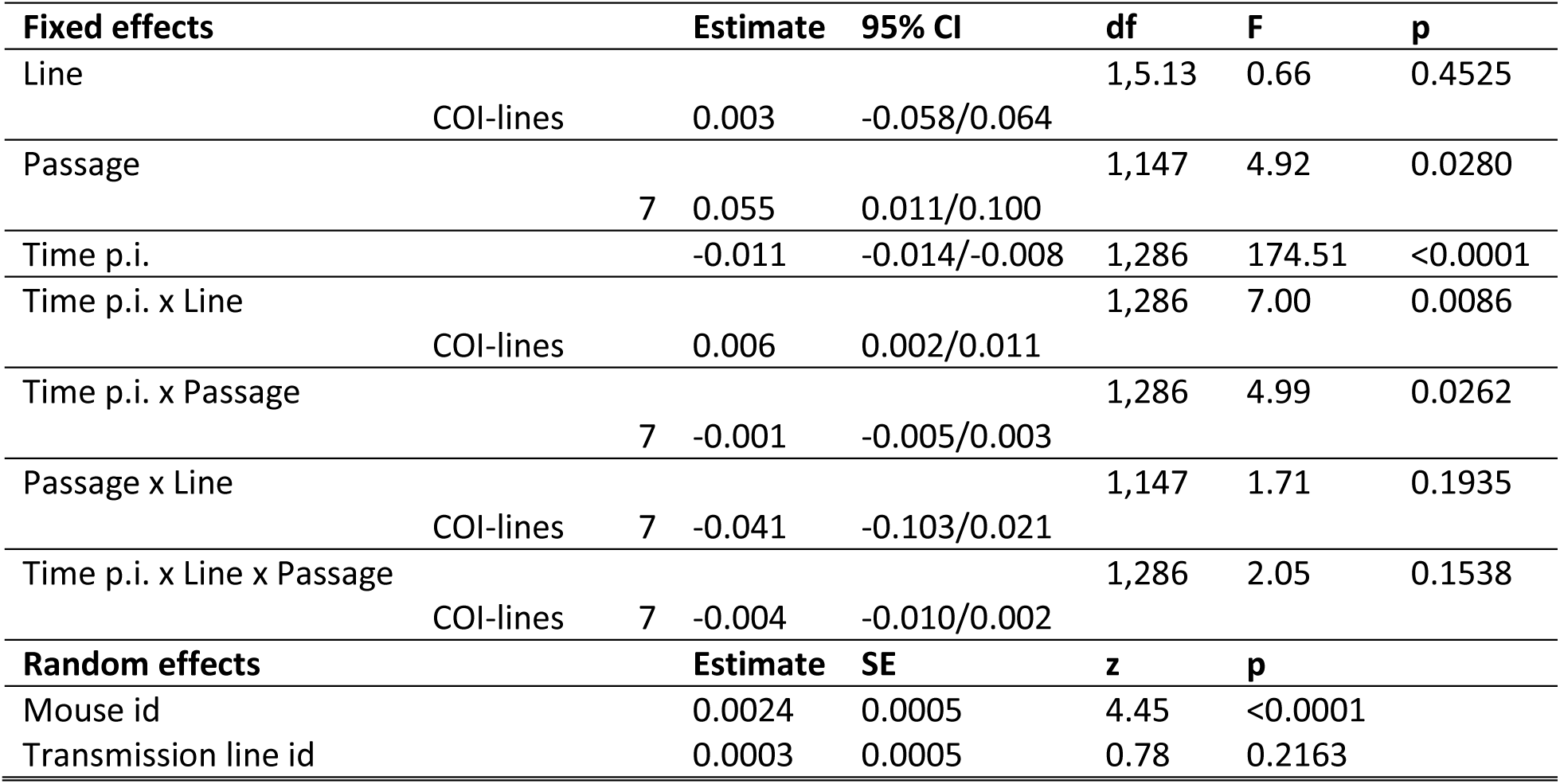
General linear mixed model with a normal distribution of errors investigating the changes in body mass over the course of the infection (variation with respect to mice infected with the ancestral Py population) in mice infected with SI-lines or COI-lines after 5 or 7 passages. For the fixed effects [time p.i., line (reference group = SI-lines), and passage (reference group = 5)], we report the parameter estimates with the 95% confidence intervals (CI), degrees of freedom (df), F and p values. Mouse identity was included as a random effect to take into account the non-independence of observations for the same individual over time; the transmission line identity was also included as a random effect to account for any heterogeneity among the replicated Py lines. For the random effects, we report the parameter estimates with the standard errors (SE), the z and p values. N = 7 transmission lines, 90 individuals and 352 observations.

The asexual reproduction of *Plasmodium* within the vertebrate host causes anemia due to the lysis of RBCs. As for changes in body mass, mice infected with the evolved Py lines lost significantly more RBCs compared to mice infected with the ancestral Py population, as shown by the highly significant effect of time p.i. in the GLMM (table 2, figure 3a,b). An intercept only GLMM that included data at day 14 p.i. showed that values were significantly lower than 1, as shown by the 95% confidence intervals non overlapping 1 [intercept (95% CI) = 0.683 (0.605/0.762), n = 48]. However, the reduction in RBCs was similar for the two lines (SI-lines and COI-lines) or across passages (table 2, figure 3a,b).

**Figure 3.**
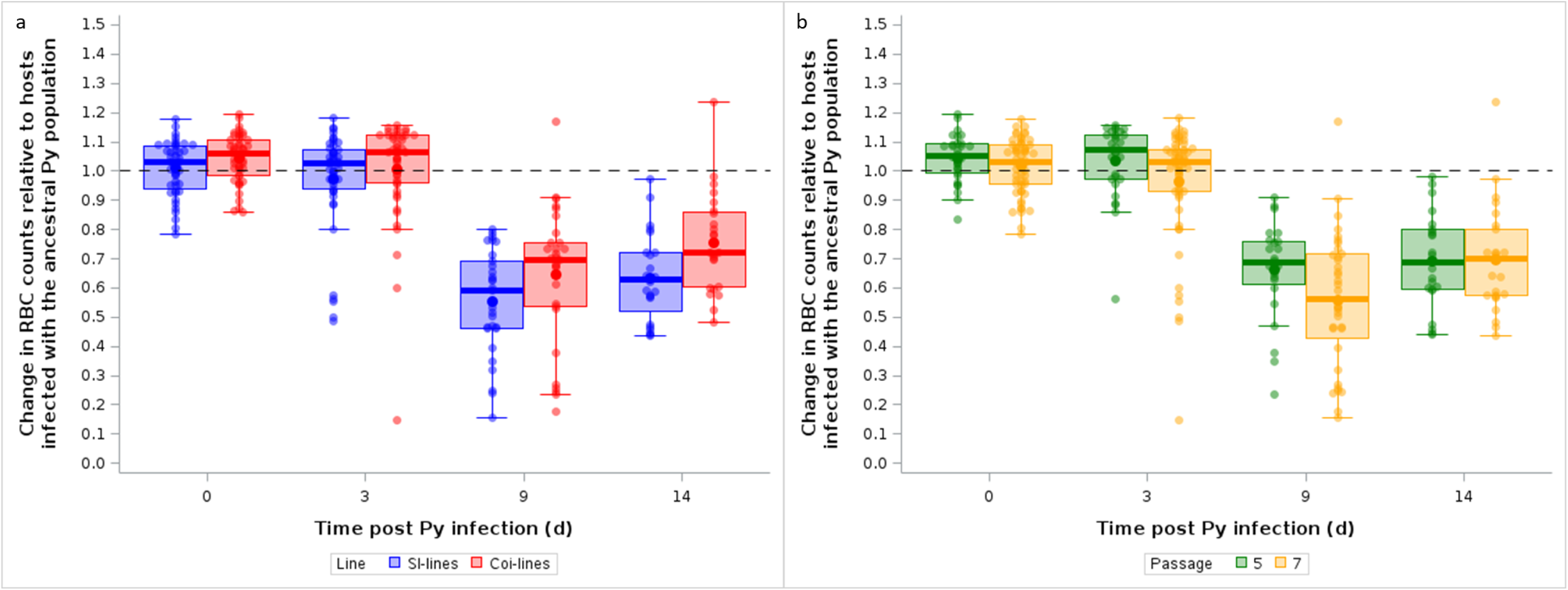
Changes in RBC counts over the course of the infection during single infection trials. RBC counts are expressed as changes relative to hosts infected with the ancestral Py population; a) Changes in RBC counts for hosts infected with SI-lines vs. COI-lines; b) Changes in RBC counts for hosts infected with Py after 5 vs. 7 passages. The dotted line represents no change with respect to hosts infected with the ancestral Py population. Dots represent the raw data, the boxes represent the interquartile range (IQR), the horizontal lines the median, and whiskers the range of data within 1.5 the IQR.

**Table 2.**
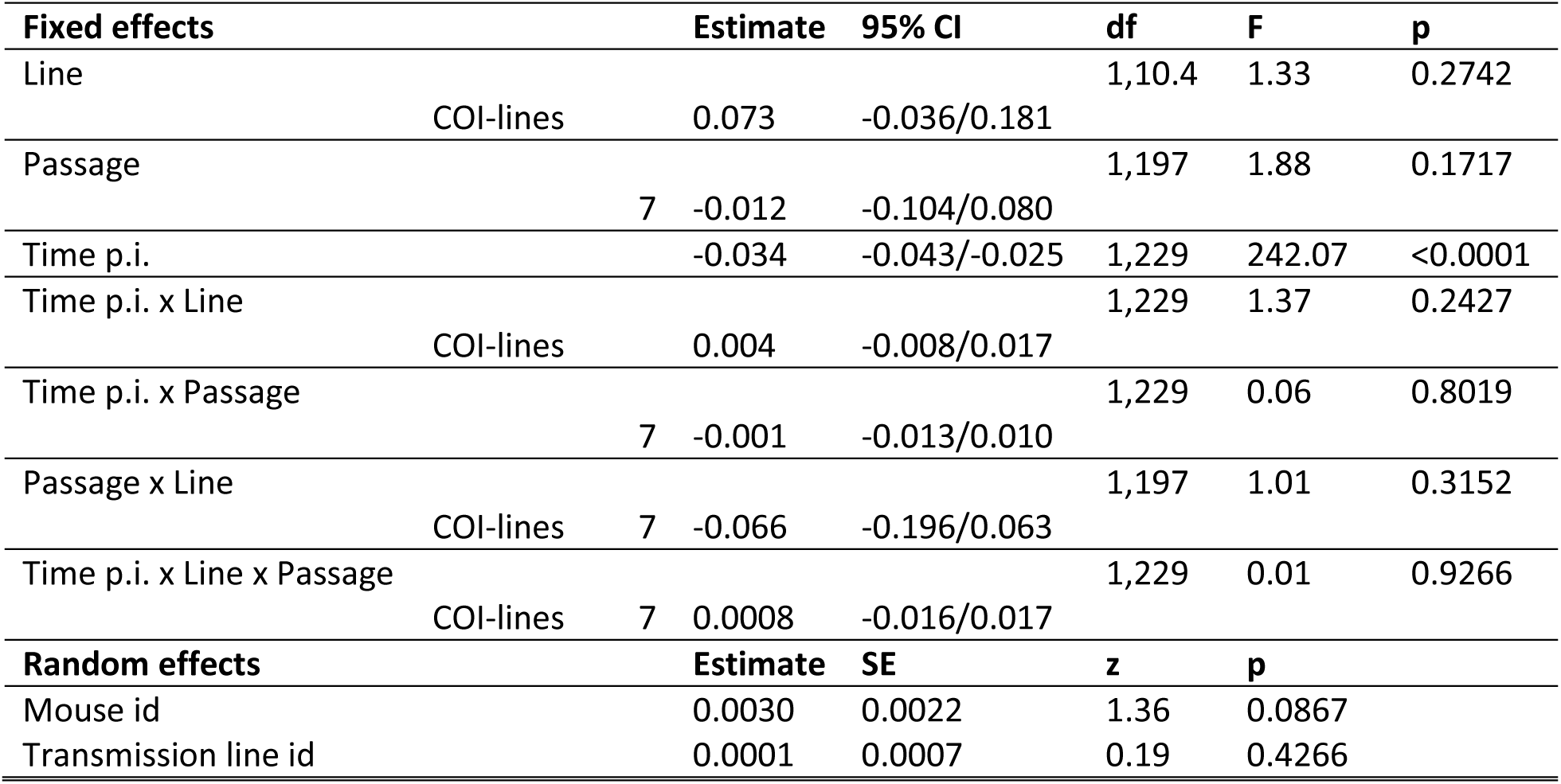
General linear mixed model with a normal distribution of errors investigating the changes in red blood cell counts over the course of the infection (variation with respect to mice infected with the ancestral Py population) in mice infected with SI-lines or COI-lines after 5 or 7 passages. For the fixed effects [time p.i., line (reference group = SI-lines), and passage (reference group = 5)], we report the parameter estimates with the 95% confidence intervals (CI), degrees of freedom (df), F and p values. Mouse identity was included as a random effect to take into account the non-independence of observations for the same individual over time; the transmission line identity was also included as a random effect to account for any heterogeneity among the replicated Py lines. For the random effects, we report the parameter estimates with the standard errors (SE), the z and p values. N = 7 transmission lines, 90 individuals and 287 observations.

#### b) Infection dynamics in hosts infected with the evolved Py lines

Serial passages are supposed to promote the spread of parasite genotypes with the fastest replication rate. In agreement with this prediction, we found that parasitemia, assessed at day 14 p.i., increased during the passages, and the rate of increase was higher for lines passaged in coinfected hosts compared to single infected hosts, as shown by the interaction between passage and line (table 3, figure 4a). However, this result does not necessarily show that parasitemia increased faster for Py lines evolving in coinfected hosts because here the two groups differed both in terms of the selective history and the current environment (single infection vs coinfection). We therefore used these evolved lines to infect mice in single infection trials, as to isolate the effect of the selection history. The model exploring the effect of lines on parasitemia during single infection trials showed that mice infected with the evolved Py lines had much higher parasitemia at day 7 and 9 p.i., compared to mice infected with the ancestral Py population (highly significant squared time term in the GLMM, table 4). The finding that values at day 14 p.i. tended to converge towards 1 shows that evolved lines induced higher parasitemia early on during the infection and then the difference plateaued (parasitemia being bounded between 0 and 1); nevertheless, parasitemia was still higher for the evolved lines compared to the ancestral population [intercept only GLMM, intercept (95% CI) = 1.139 (1.090/1.188), n = 48; figure 4b,c)]. The model also suggested that the relationship between changes in parasitemia and time was more concave after seven passages (higher peak at day 7 p.i.) (table 4), but visual inspection of figure 4c showed that the effect size was small. We did not find any difference between SI-lines and COI-lines (table 4, figure 4b).

**Figure 4.**
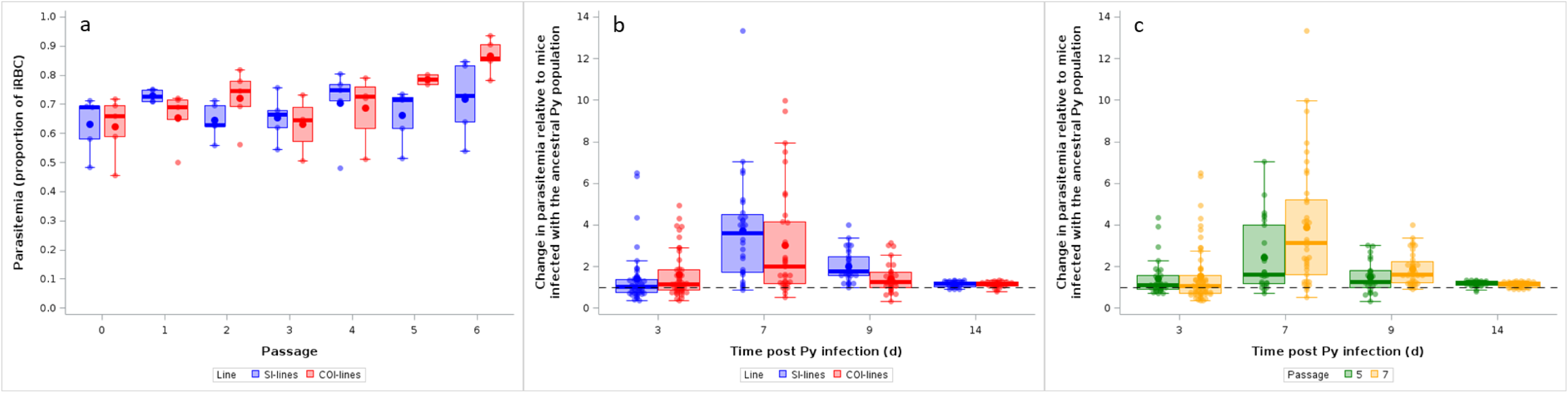
a) Parasitemia (proportion of iRBC) at day 14 p.i. during the passages in single infected or coinfected hosts; b) Changes in parasitemia over the course of the infection in hosts infected with SI-lines vs. COI-lines during single infection trials. Parasitemia is expressed as change relative to hosts infected with the ancestral Py population; c) Changes in parasitemia over the course of the infection in hosts infected after 5 vs. 7 passages during single infection trials. Parasitemia is expressed as change relative to hosts infected with the ancestral Py population. The dotted line represents no change with respect to hosts infected with the ancestral Py population. For the three panels, dots represent the raw data, the boxes represent the interquartile range (IQR), the horizontal lines the median, and whiskers the range of data within 1.5 the IQR.

**Table 3.**
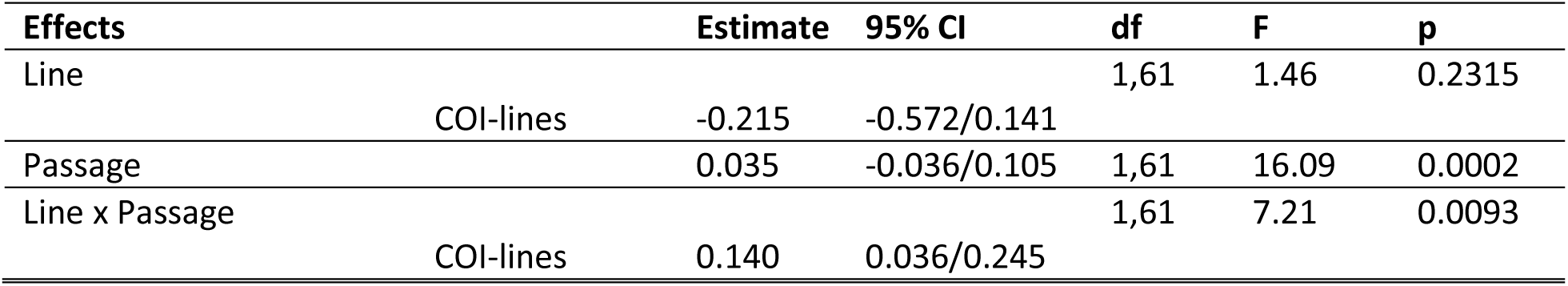
Generalized linear model with a beta-distribution of errors investigating the changes in parasitemia (proportion of iRBCs) at day 14 p.i. during the passages in single infected or coinfected hosts. Passage, line (reference = SI lines) and the two-way interaction were included independent variables, for which we report the parameter estimates with the 95% confidence intervals (CI), degrees of freedom (df), F and p values. N = 65 individuals.

**Table 4.**
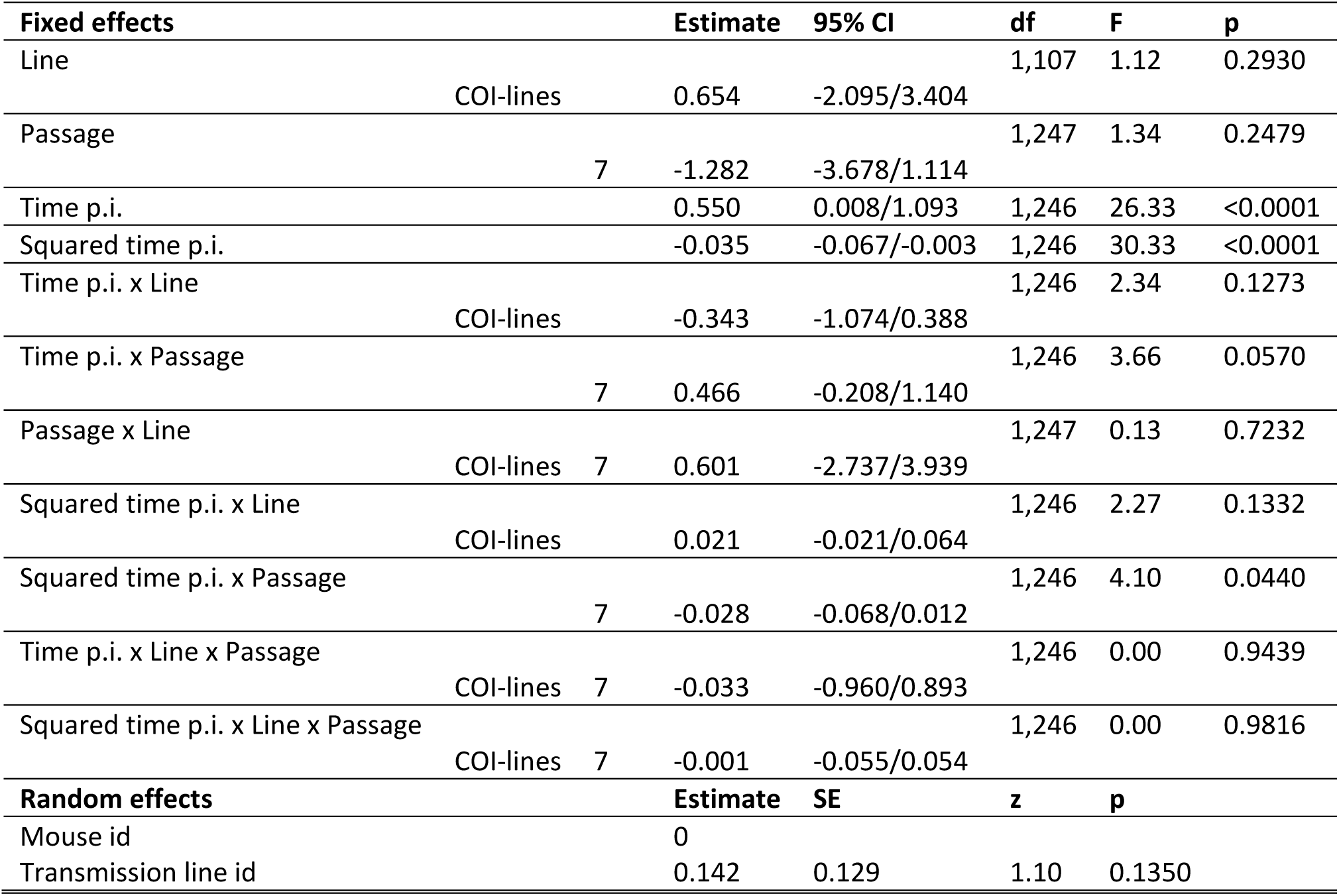
General linear mixed model with a normal distribution of errors investigating the changes in parasitemia over the course of the infection (variation with respect to mice infected with the ancestral Py population) in mice infected with SI-lines or COI-lines after 5 or 7 passages. For the fixed effects [time p.i., squared time p.i., line (reference group = SI-lines), and passage (reference group = 5)], we report the parameter estimates with the 95% confidence intervals (CI), degrees of freedom (df), F and p values. Mouse identity was included as a random effect to take into account the non-independence of observations for the same individual over time; the transmission line identity was also included as a random effect to account for any heterogeneity among the replicated Py lines. For the random effects, we report the parameter estimates with the standard errors (SE), the z and p values. N = 7 transmission lines, 90 individuals and 261 observations.

#### c) Tolerance to the infection in hosts infected with the evolved Py lines

Severity of disease symptoms does not depend only on the damage caused by parasite replication, but also on the capacity of the host to repair it (Medzhitov et al. 2012). As mentioned above, *Plasmodium* replication causes the lysis of RBCs and induces anemia. Lost RBCs can, however, be replaced by hematopoiesis, reducing the net cost of the infection. We therefore assessed whether mice infected with the evolved lines (after seven passages) differed in the amount of circulating erythropoietin, the hormone regulating the production of RBCs, at day 14 post Py infection. An intercept only GLMM showed that there was no difference between the ancestral and the evolved lines [intercept (95% CI) = 1.900 (0.772/3.027)]. Similarly, there was no difference between the evolved lines (SI-lines and COI-lines) (GLMM, F_1,15_ = 0.29, p = 0.5974; figure 5a).

**Figure 5.**
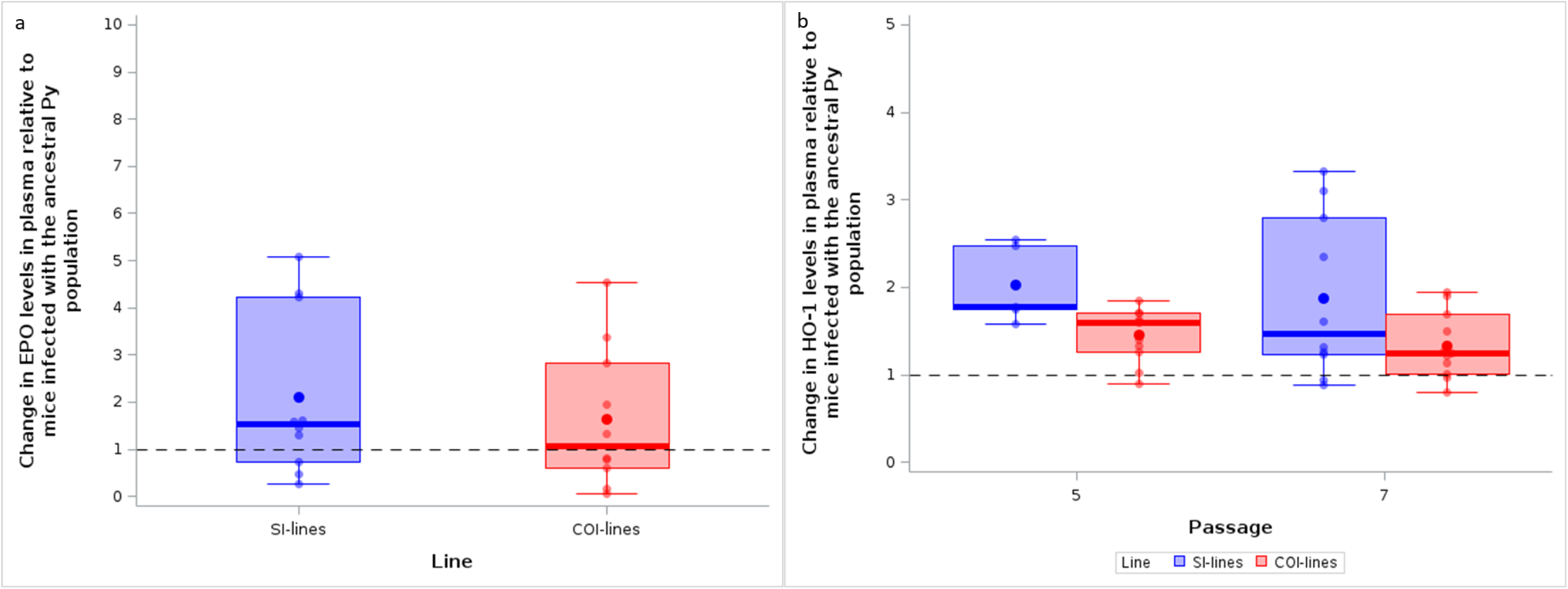
a) Plasma level of EPO at day 14 p.i. in hosts infected with SI-lines vs. COI-lines after 7 passages during single infection trials. EPO is expressed as change relative to hosts infected with the ancestral Py population; b) Plasma levels of HO-1 at day 14 p.i. in hosts infected with SI-lines vs. COI-lines after 5 and 7 passages during single infection trials. HO-1 is expressed as change relative to hosts infected with the ancestral Py population. The dotted line represents no change with respect to hosts infected with the ancestral Py population. Dots represent the raw data, the boxes represent the interquartile range (IQR), the horizontal lines the median, and whiskers the range of data within 1.5 the IQR.

Lysis of RBCs releases free heme groups that have strong pro-oxidant activities and have been shown to be an important component of *Plasmodium* pathogenesis (Kumar and Bandyopadhyay 2005, Seixas et al. 2009). We therefore investigated whether evolved Py lines elicited a different response in terms of the production of the scavenging enzyme heme oxygenase-1 (HO-1) at day 14 post-infection. We found that, overall, hosts infected with the evolved lines had a much stronger response in terms of HO-1, compared to mice infected with the ancestral Py population [intercept only GLMM, intercept (95% CI) = 1.731 (1.320/2.142), n = 37]. The GLMM comparing the evolved lines showed that COI-lines induced a weaker HO-1 response in the hosts compared to SI-lines (table 5; figure 5b).

**Table 5.**
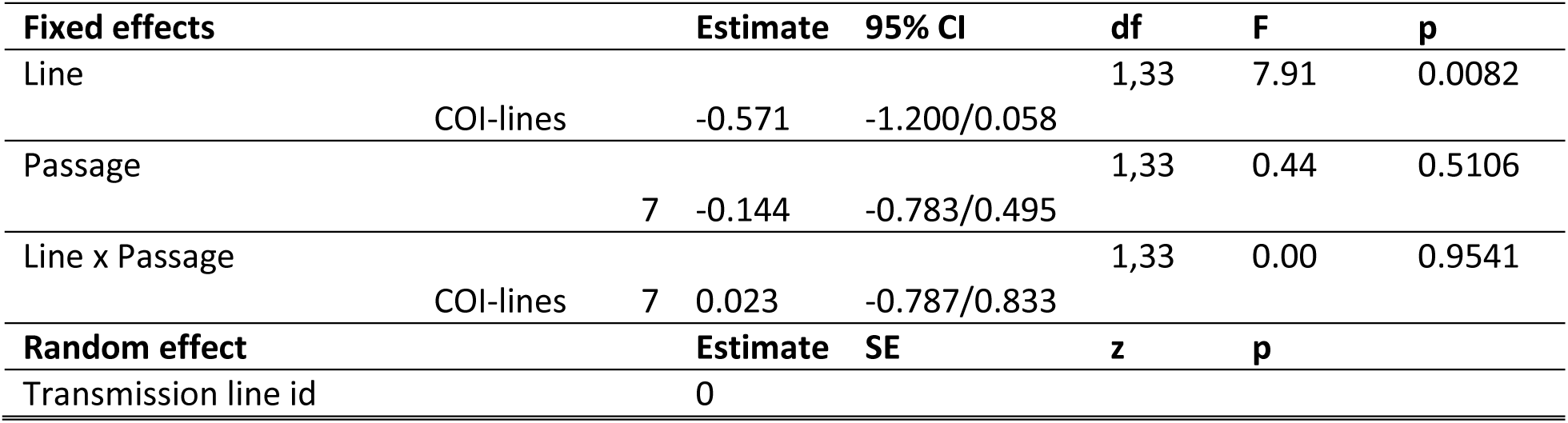
General linear mixed model with a normal distribution of errors investigating the changes in HO-1 levels in plasma at day 14 p.i. (variation with respect to mice infected with the ancestral Py population) in mice infected with SI-lines or COI-lines after 5 or 7 passages. For the fixed effects [line (reference group = SI-lines), and passage (reference group = 5)], we report the parameter estimates with the 95% confidence intervals (CI), degrees of freedom (df), F and p values. The transmission line identity was also included as a random effect to account for any heterogeneity among the replicated Py lines. Mouse identity was not included in the model, because each mouse provided a single HO-1 measurement. For the random effect, we report the parameter estimates with the standard errors (SE), the z and p values. N = 7 lines and 37 observations.

Whatever the possible mechanisms underlying the host capacity to minimize the cost of the infection, a more direct test of the difference in tolerance between groups is to regress a proxy of the cost of infection on parasitemia (Råberg et al. 2007); the steeper the slope of the regression, the weaker the tolerance. We found that mice infected with the evolved lines after seven passages were less tolerant to Py infection (steeper slope between RBC counts and parasitemia) compared to mice infected with the ancestral population (table 6; figure 6a,b). However, tolerance did not differ between mice infected with SI-lines or COI-lines (figure 6c,d).

**Figure 6.**
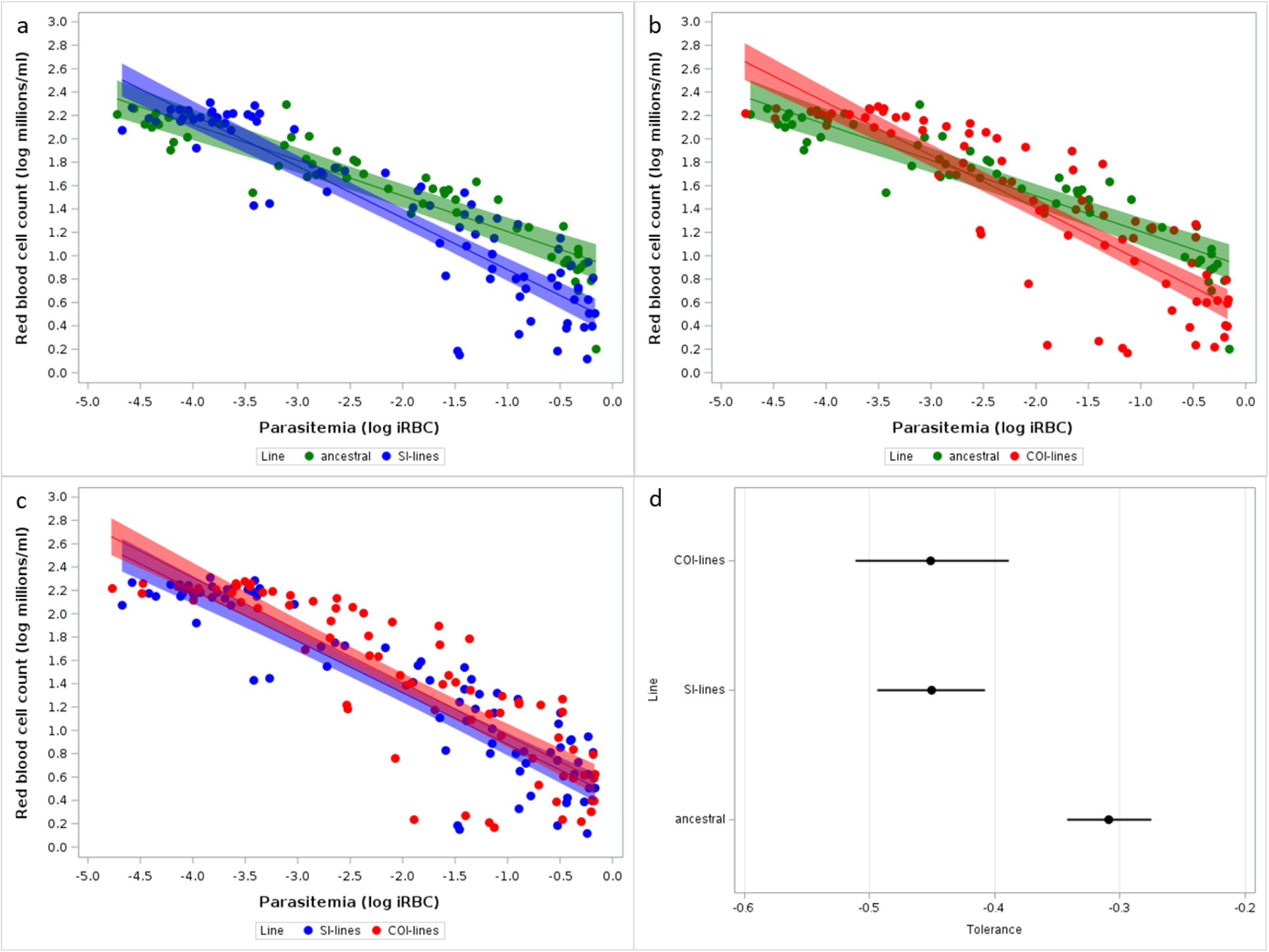
Tolerance to the infection in hosts infected with the ancestral and the evolved Py lines. To improve readability, the three treatment groups were split to allow pair-wise comparisons of a) ancestral vs SI-lines; b) ancestral vs COI-lines; c) SI-lines vs COI-lines, although the model was run on the three groups (see text). Tolerance is expressed as the slope of the regression of RBC count on parasitemia (proportion of iRBC) (log-log plot); d) Forest plot showing the estimates of the slopes with the 95% confidence intervals for each of the three groups.

**Table 6.** General linear mixed model with a normal distribution of errors investigating the relationship between red blood cell counts and parasitemia in mice infected with Py from the ancestral population, infected with SI-lines or infected with COI-lines. For the fixed effects [line (reference group = ancestral), and parasitemia], we report the parameter estimates with the 95% confidence intervals (CI), degrees of freedom (df), F and p values. Mouse identity was included as a random effect to take into account the non-independence of observations for the same individual. For the random effect, we report the parameter estimates with the standard errors (SE), the z and p values. N = 75 individuals and 220 observations.

| **Fixed effects** |  | **Estimate** | **95% CI** | **df** | **F** | **p** |
| --- | --- | --- | --- | --- | --- | --- |
| Line |  |  |  | 1,142 | 765.78 | <0.0001 |
|  | SI-lines | -0.462 | -0.662/-0.262 |  |  |  |
|  | COI-lines | -0.396 | -0.601/-0.190 |  |  |  |
| Parasitemia |  | -0.305 | -0.357/-0.253 | 2,142 | 11.39 | <0.0001 |
| Line x Parasitemia |  |  |  | 2,142 | 10.16 | <0.0001 |
|  | SI-lines | -0.136 | -0.205/-0.067 |  |  |  |
|  | COI-lines | -0.146 | -0.219/-0.074 |  |  |  |
| **Random effect** |  | **Estimate** | **SE** | **z** | **p** |  |
| Mouse ID |  | 0.017 | 0.009 | 1.90 | 0.0287 |  |

#### d) Virulence of the evolved Py lines

During the passages, Py was transmitted from mouse to mouse at day 14 p.i. (approximately the day of peak parasitemia), and due to host mortality occurring before transmission, some of the Py lines were lost. Pre-transmission host mortality was higher for the coinfection passages (Log-rank test, χ²_1_ = 5.46, p = 0.0195, figure 7a) and, up to passage 7, more lines were lost in the coinfection passages (3 out of 5) than in the single infection passages (1 out of 5). When tested in single infection trials, we found that both evolved lines elicited higher host mortality than the ancestral population (Log-Rank test, χ²_2_ = 31.22, p < 0.0001, figure 7b; both pairwise comparisons, Sidak adjusted p’s < 0.0001); however, there was no difference between the SI-lines and the COI-lines (Sidak adjusted p = 0.6688, figure 7b).

**Figure 7.**
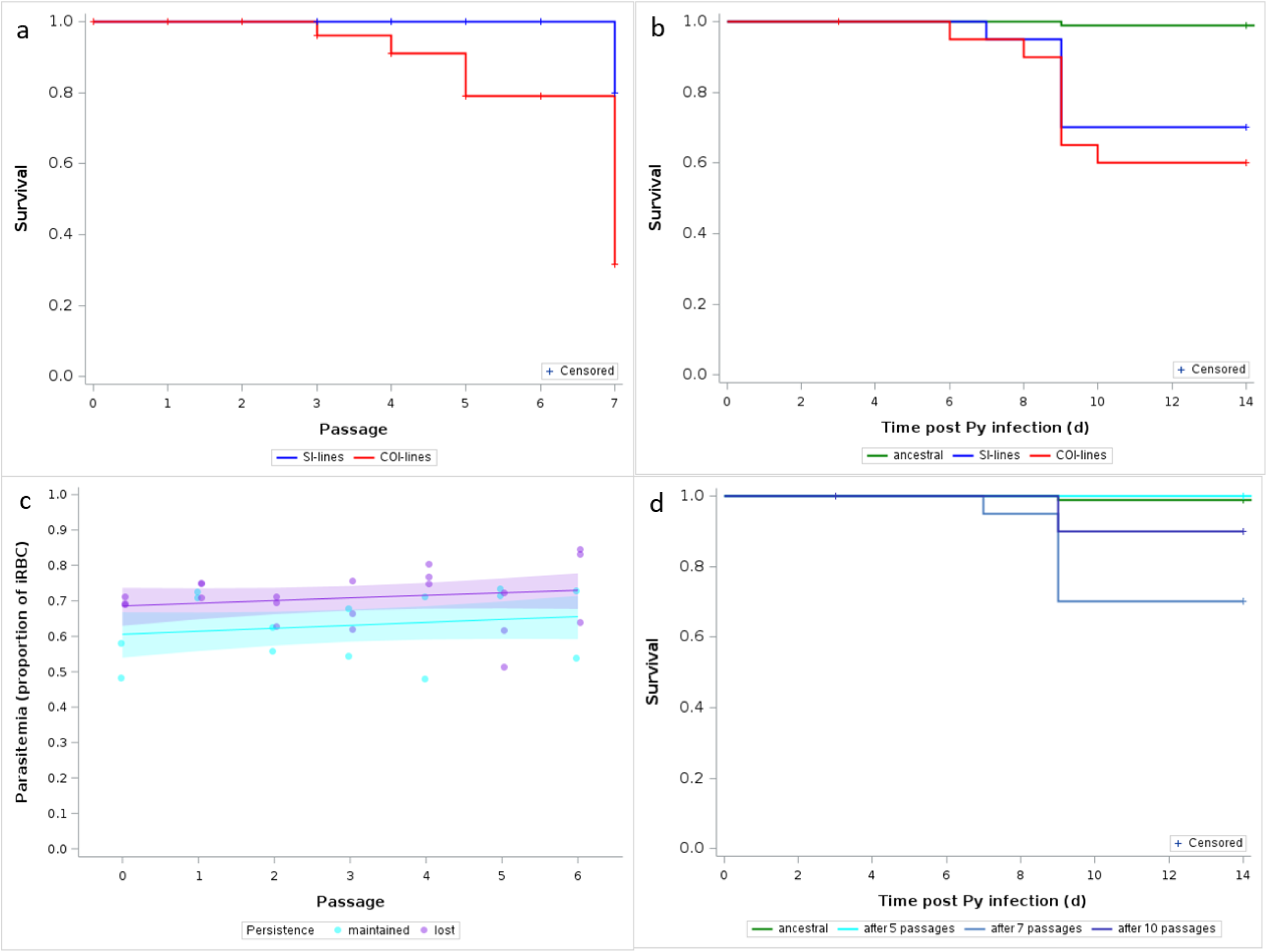
a) Survival curves of single infected and coinfected hosts during the passages; b) Survival over the course of the infection of hosts infected after 7 passages; c) Parasitemia (proportion of iRBC) at day 14 p.i. of single infected hosts during the passages, according to the fate of the lines (blue = lines that persisted up to passage 10; red = lines that were lost). Dots represent the raw data, the lines the model fit and the shaded area the 95% CI around the fit; d) Survival over the course of the infection of hosts infected with the ancestral Py population or with SI-lines after 5, 7 and 10 passages. Crosses indicate censored observations.

For the single infection passages, Py was transmitted up to passage 10. This allowed us to check whether the cost of virulence (pre-transmission host mortality) also occurred in this group over a higher number of passages. Indeed, the number of Py lines lost kept increasing up to passage 10 (60% lines lost), as did mortality (figure S2). In agreement with the hypothesis that the lines with the fastest replication were the most virulent and more likely to be lost during the passages, we found that at passage 6 (before any line had been lost) the parasitemia at day 14 p.i. was higher for lines that were subsequently lost (F_1,32_ = 7.82, p = 0.0087, figure 7c). We also found that after ten passages, the persisting lines that were used for the evaluation trials had lower virulence compared to the lines tested after seven passages (Sidak adjusted p = 0.0055; figure 7d), but still higher than the ancestral population (Sidak adjusted p = 0.0363; figure 7d). Therefore, by transmitting Py at day 14 p.i., we allowed selection to operate against the most virulent lines.

#### e) Immune response of hosts infected with the evolved Py lines

Infection with *Plasmodium* induces a strong pro-inflammatory response which, in turn, induces the production of regulatory mechanisms as to resolve inflammation. We therefore wished to test whether evolved Py lines elicited different pro- and anti-inflammatory responses in mice compared to the ancestral Py population. To this aim, we assessed the gene expression of the pro-inflammatory cytokine IFN-γ and the anti-inflammatory cytokine IL-10 in splenocytes at day 3 and 14 post Py infection. Overall, the gene expression (ΔΔ Cts) of the evolved lines did not differ from the ancestral population, as shown by an intercept only GLMM [IFN-γ, intercept (95% CI) = -0.471 (-1.235/0.293); IL-10, intercept (95% CI) = -0.172 (-1.031/0.687)]. However, after seven passages, SI-lines elicited an upregulated expression of IFN-γ compared to COI-lines at day 3 p.i., but this difference vanished by day 14 p.i. (table 7, figure 8a). This result was mirrored by the findings on the gene expression of the regulatory cytokine IL-10. SI-lines tended to elicit an upregulated expression of IL-10 compared to COI-lines at day 3 p.i., and at day 14 p.i. the two groups had identical ΔΔ Ct values (table 8, figure 8b). Finally, we assessed whether the evolved lines elicited a different expansion of regulatory T cells in the spleen at day 14 p.i. and found that overall hosts infected with the evolved lines had a lower proportion of FoxP3^+^ cells among CD4^+^ lymphocytes [intercept only GLMM, intercept (95% CI) = 0.884 (0.777/0.992), n = 43]. The comparison between evolved lines indicated that that COI-lines elicited a stronger expansion of the population of Treg cells compared to the SI-lines (table 9, figure 8c).

**Figure 8.**
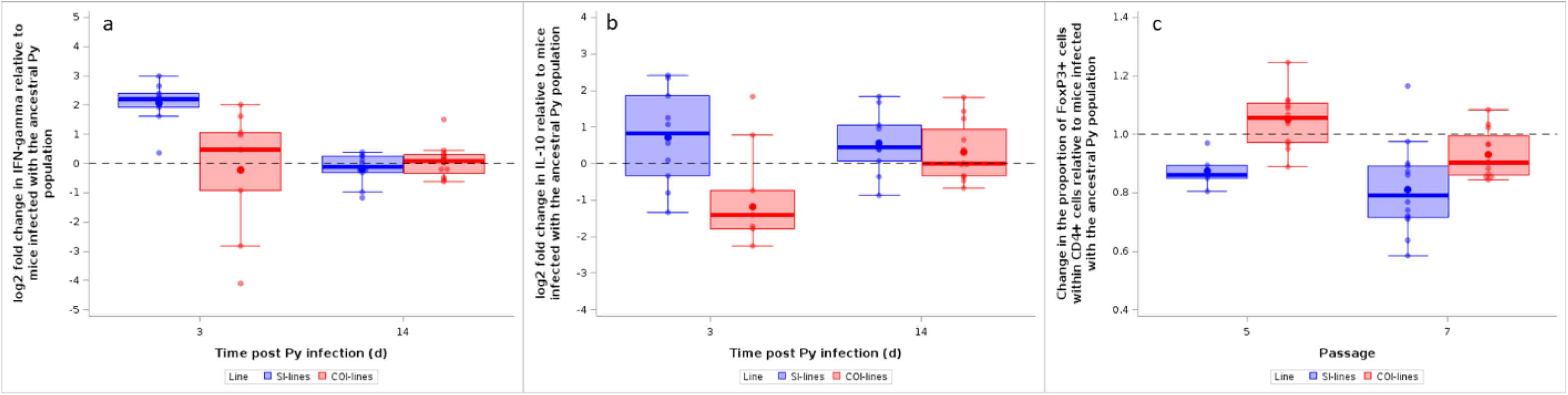
a) log_2_ fold change of *IFN-γ* mRNA in spleen at day 3 and 14 p.i. in hosts infected with SI-lines or COI-lines after 7 passages; b) log_2_ fold change of *IL-10* mRNA in spleen at day 3 and 14 p.i. in hosts infected with SI-lines or COI-lines; c) Change in the proportion of FoxP3^+^ cells within CD4^+^ T cells at day 14 p.i. in hosts infected with SI-lines or COI-lines after 5 and 7 passages. The dotted line represents no change with respect to hosts infected with the ancestral Py population. Dots represent the raw data, the boxes represent the interquartile range (IQR), the horizontal lines the median, and whiskers the range of data within 1.5 the IQR.

**Table 7.** General linear mixed model with a normal distribution of errors investigating IFN-γ gene expression (ΔΔ Ct with respect to mice infected with the ancestral Py population) at day 3 and 14 p.i. in mice infected with SI-lines or COI-lines after 7 passages. For the fixed effects [time p.i. (reference = 3), and line (reference group = SI-lines)], we report the parameter estimates with the 95% confidence intervals (CI), degrees of freedom (df), F and p values. The transmission line identity was included as a random effect to account for any heterogeneity among the replicated Py lines. Mouse identity was not included in the model, because each mouse provided a single ΔΔ Ct measurement. For the random effect, we report the parameter estimates with the standard errors (SE), the z and p values. N = 5 lines and 41 observations.

| **Fixed effects** |  |  | **Estimate** | **95% CI** | **df** | **F** | **p** |
| --- | --- | --- | --- | --- | --- | --- | --- |
| Line |  |  |  |  | 1,37 | 8.59 | 0.0058 |
|  | COI-lines |  | 2.285 | 1.260/3.309 |  |  |  |
| Time p.i. |  |  |  |  | 1,37 | 8.17 | 0.0070 |
|  |  | 14 | 2.259 | 1.262/3.257 |  |  |  |
| Time p.i. x Line |  |  |  |  | 1,37 | 13.55 | 0.0007 |
|  | COI-lines | 14 | -2.544 | -3.944/-1.144 |  |  |  |
| **Random effect** |  |  | **Estimate** | **SE** | **z** | **p** |  |
| Transmission line id |  |  | 0 |  |  |  |  |

**Table 8.**
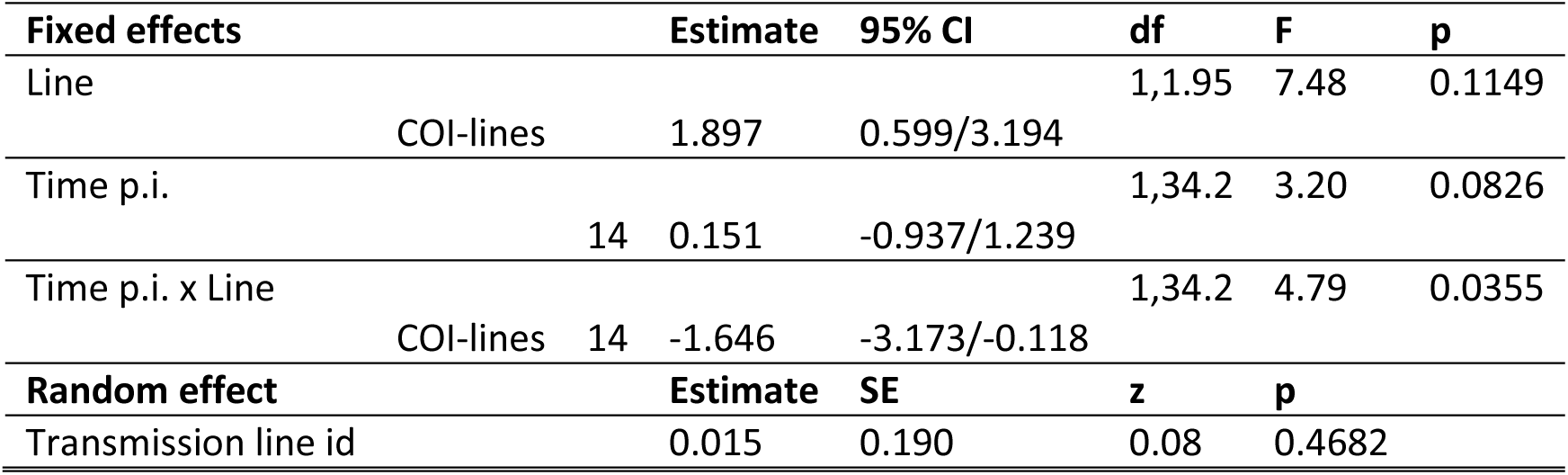
General linear mixed model with a normal distribution of errors investigating IL-10 gene expression (ΔΔ Ct with respect to mice infected with the ancestral Py population) at day 3 and 14 p.i. in mice infected with SI-lines or COI-lines after 7 passages. For the fixed effects [time p.i. (reference = 3), and line (reference group = SI-lines)], we report the parameter estimates with the 95% confidence intervals (CI), degrees of freedom (df), F and p values. The transmission line identity was included as a random effect to account for any heterogeneity among the replicated Py lines. Mouse identity was not included in the model, because each mouse provided a single ΔΔ Ct measurement. For the random effect, we report the parameter estimates with the standard errors (SE), the z and p values. N = 5 lines and 41 observations.

**Table 9.**
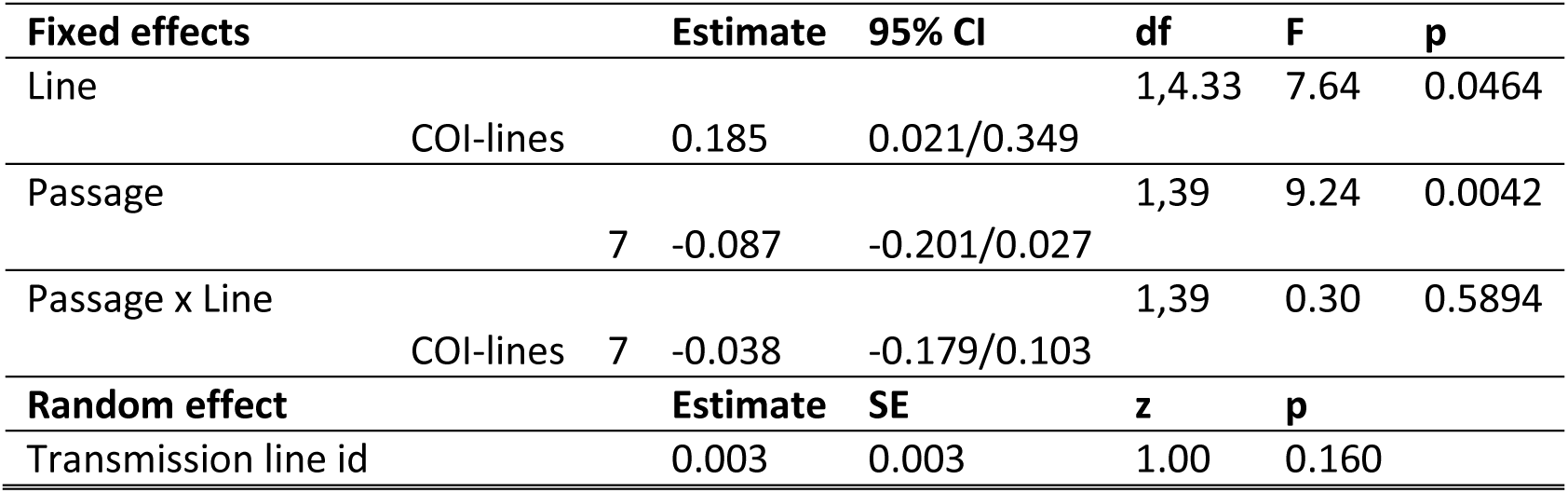
General linear mixed model with a normal distribution of errors investigating the changes in the proportion of FoxP3^+^ cells within CD4^+^ lymphocytes (variation with respect to mice infected with the ancestral Py population) at day 14 p.i. in mice infected with SI-lines or COI-lines after 7 passages. For the fixed effects [passage (reference = 5), and line (reference group = SI-lines)], we report the parameter estimates with the 95% confidence intervals (CI), degrees of freedom (df), F and p values. The transmission line identity was included as a random effect to account for any heterogeneity among the replicated Py lines. Mouse identity was not included in the model, because each mouse provided a single measurement of the proportion of FoxP3^+^ cells. For the random effect, we report the parameter estimates with the standard errors (SE), the z and p values. N = 7 lines and 43 observations.

### 2) Effect of mismatched environment on *Plasmodium* adaptation

In a second experiment, we evaluated COI-lines in coinfected hosts, which allowed us to investigate whether Py performance changed according to the mismatch between the environment encountered during the passages and the environment faced during the evaluation trials. We had two matched groups (SI-lines tested in si-hosts, COI-lines tested in coi-hosts) and one mismatched group (COI-lines tested in si-hosts) (capital letters referring to the passage treatment and the small letters the evaluation trial treatment).

We used parasite replication as a proxy of parasite performance and found that the infection dynamics differed between the three groups as shown by the time x trial and squared time x trial interactions (table S3, figure S3). In agreement with the prediction, the mismatched group (COI/si) had lower parasitemia compared to the other groups at day 7 and 9 p.i., with parasitemia converging towards high values at day 14 p.i. (figure S3). This finding was corroborated by an additional model where the two matched environment groups were clustered together and tested against the mismatched group, showing a slower replication rate of Py in mismatched environments (table 10, figure 9). However, there was no difference in infection-induced host mortality between matched- and mismatched-environment groups (Log-rank Test, χ²1 = 0.002, p = 0.9656; figure S4), and hosts had similar tolerance when infected with matched- or mismatched-environment Py (GLMM, parasitemia x matched environment, F_1,178_ = 0.10, p = 0.7560, n = 85 individuals and 216 observations; figure S5).

**Figure 9.**
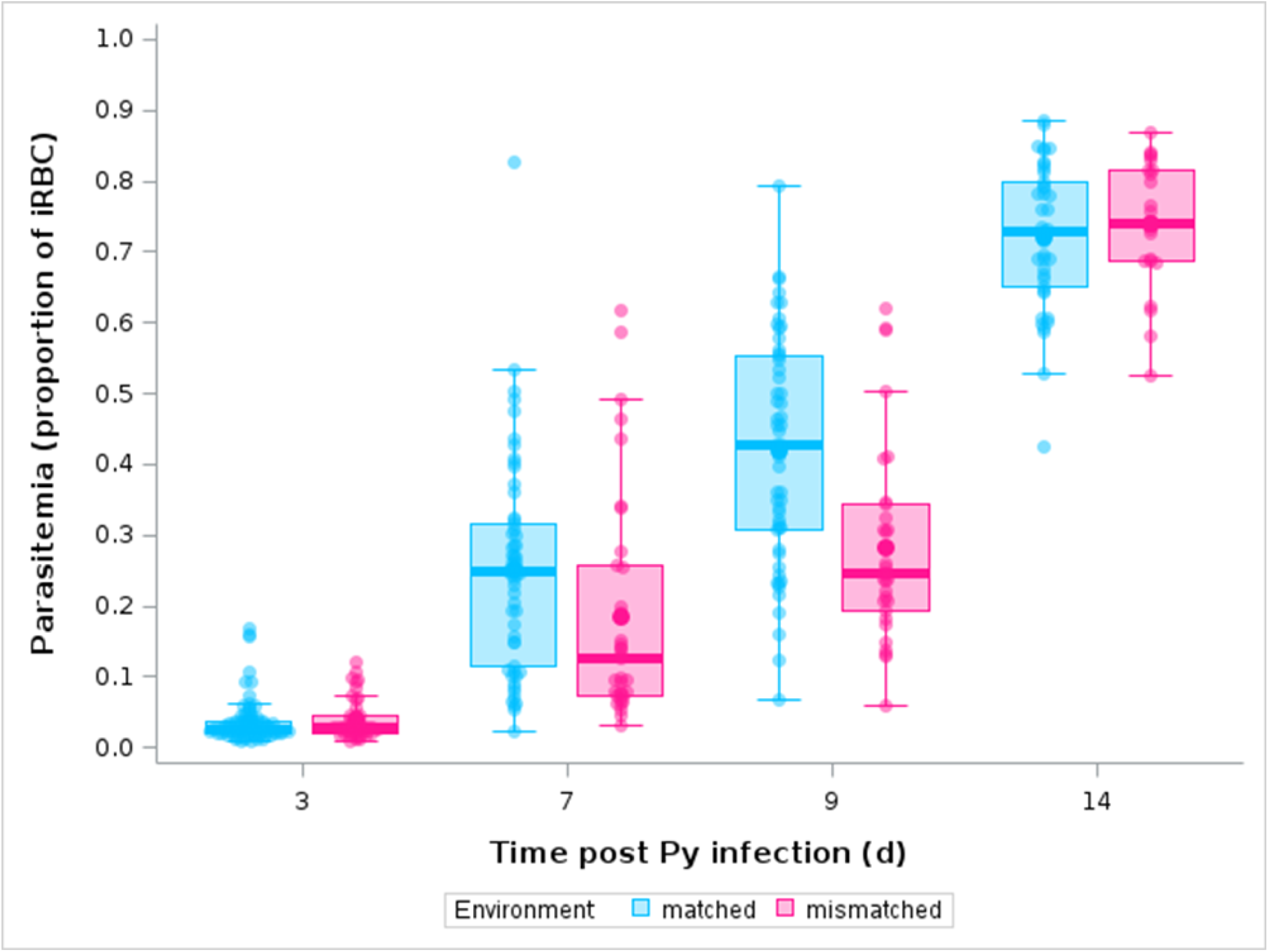
Parasitemia over the course of the infection in hosts infected with Py that experienced matched environments between the passages and the evaluation trials or with Py that experienced mismatched environments. Dots represent the raw data, the boxes represent the interquartile range (IQR), the horizontal lines the median, and whiskers the range of data within 1.5 the IQR.

**Table 10.** Generalized linear mixed model with a beta-distribution of errors investigating the changes in parasitemia over the course of the infection in mice infected with Py having encountered matched or mismatched environments between the passages and the evaluation trials. Trials were conducted after 5 and 7 passages. For the fixed effects [time p.i., squared time p.i., passage (reference = 5), and environment (reference group = matched)], we report the parameter estimates with the 95% confidence intervals (CI), degrees of freedom (df), F and p values. Mouse identity was included as a random effect to take into account the non-independence of observations for the same individual over time; the transmission line identity was also included as a random effect to account for any heterogeneity among the replicated Py lines. For the random effects, we report the parameter estimates with the standard errors (SE), the z and p values. N = 130 individuals, 7 lines and 373 observations.

| Fixed effects |  |  | Estimate | 95% CI | df | F | p |
| --- | --- | --- | --- | --- | --- | --- | --- |
| Time p.i. |  |  | 0.575 | 0.403/0.746 | 1,274 | 138.87 | <0.0001 |
| Squared time p.i. |  |  | -0.010 | -0.019/-0.001 | 1,271.8 | 13.63 | 0.0003 |
| Passage |  |  |  |  | 1,291 | 1.46 | 0.2280 |
|  | 7 |  | -0.822 | -1.828/0.183 |  |  |  |
| Environment |  |  |  |  | 1,297.1 | 3.44 | 0.0647 |
|  |  | mismatched | 0.524 | -0.902/1.950 |  |  |  |
| Time p.i. x Passage |  |  |  |  | 1,274 | 7.90 | 0.0053 |
|  | 7 |  | 0.341 | 0.120/0.562 |  |  |  |
| Time p.i. x Environment |  |  |  |  | 1,273.9 | 13.89 | 0.0002 |
|  |  | mismatched | -0.287 | -0.593/0.019 |  |  |  |
| Environment x Passage |  |  |  |  | 1,290.3 | 0.45 | 0.5037 |
|  | 7 | mismatched | 0.586 | -1.137/2.309 |  |  |  |
| Squared time p.i. x Passage |  |  |  |  | 1,272 | 10.49 | 0.0013 |
|  | 7 |  | -0.020 | -0.031/-0.008 |  |  |  |
| Squared time p.i. Environment |  |  |  |  | 1,271.83 | 18.86 | <0.0001 |
|  |  | mismatched | 0.018 | 0.002/0.034 |  |  |  |
| Time p.i. x Passage x Environment |  |  |  |  | 1,273.9 | 0.54 | 0.4616 |
|  | 7 | mismatched | -0.142 | -0.520/0.237 |  |  |  |
| Squared time p.i. x Passage x Environment |  |  |  |  | 1,271.8 | 0.51 | 0.4739 |
|  | 7 | mismatched | 0.007 | -0.012/0.027 |  |  |  |
| Random effects |  |  | Estimate | SE | z | p |  |
| Mouse id |  |  | 0.136 | 0.034 | 4.00 | <0.0001 |  |
| Transmission line id |  |  | 0.085 | 0.060 | 1.40 | 0.0805 |  |

## Discussion

We showed that serial passages of Py rapidly resulted in faster replication rate and increased virulence; however, SI-lines and COI-lines had similar virulence (and more generally induced disease symptoms of similar severity), and hosts similarly tolerated the infections with the two selection lines when evaluated during single infection trials. At first glance, this might be seen as evidence that environmental (within-host) conditions encountered in single infected and coinfected hosts exerted similar selection pressures on Py, but it is also possible that the cost of virulence incurred by the strains with the fastest replication rate during the passages, constrained virulence evolution. We also found evidence suggesting that Py rapidly adapted to the conditions provided by the host since it had a faster replication rate when the evaluation trials matched the environment experienced during the passages.

Parasitic microorganisms have the capacity to rapidly adapt to novel environments due to their large population size, short generation time and high mutation rate (Drew et al. 2021). Examples of parasite adaptation to novel environments (or novel host genotypes/phenotypes) are manifold and come from a variety of approaches based both on lab experiments and field observations (Mackinnon and Read 2004, Barclay et al. 2012, Kubinak and Potts 2013, Barclay et al. 2014, Rafaluk et al. 2015, Urbanowicz et al. 2016, Rono et al. 2018).

In our experimental approach, Py did not face a novel host *sensu stricto* but experienced different environmental conditions according to the coinfection status of the host. Indeed, in a previous work, we showed that when Py infects hosts previously infected with Hp, the infection dynamics is altered and virulence of Py increases compared to single infection (Dusuel et al. 2025). The mechanisms underlying the effects of coinfection are essentially related to the regulation of the anti-malarial immune response (top-down regulation), providing a favorable ground for Py replication in coinfected hosts (Dusuel et al. 2025). Given that faster replication rate is correlated to the production of transmissible stages, this result suggests higher Py fitness in coinfected hosts, unless host death occurs prior to transmission.

Therefore, the follow-up question was, does the immune environment provided by a single infected or a coinfected host select for Py with specific replication rate and virulence? To predict the direction of a possible microevolutionary response, we tentatively used the theoretical framework developed by Fenton (2013) who predicted that under some specific conditions, deworming should select for increased pathogen virulence. Although the structure of the model differs with many respects from our experimental approach, there are also several analogies between them. In particular, Fenton (2013) considered different types of within-host interactions between helminths and microparasites, both antagonistic and synergistic effects. Among these, he explicitly considered two cases that match the interaction between Py and Hp: helminths aggravate the severity of symptoms caused by microparasites, and hosts infected with helminths have a reduced recovery rate. Under these specific conditions, the model predicted that in the absence of helminth infection (e.g., during deworming), microparasite virulence should increase.

Our results did not provide a support to this prediction and, if any, tended rather to indicate that Py passaged in coinfected hosts might have acquired higher virulence compared to lines passaged in single infected mice. This is based on the findings that the lines with the fastest replication rate, inducing a pre-transmission host mortality, were lost during the passages, which occurred more in the coinfection treatment. Therefore, the subset of lines still available for the evaluation trials was probably skewed towards the most benign ones, especially in the coinfection treatment. It should be reminded that we adopted a transmission rule where Py was passaged at a set, and relatively late, time point, irrespective of the time when symptoms developed. This differs from the other strategy usually implemented in serial passage experiments where pathogens are transmitted before the onset of symptoms (at an early time point) (Rafaluk et al. 2015). Passaging pathogens early or late during the infectious period, obviously changes the selection pressures acting on replication rate and virulence. When passages occur early on during the infectious period, pathogen strains with the fastest replication rate outnumber slow replicating strains in the inoculum that is transferred between hosts. In addition to this, given that transmission occurs before host death, fast replicating strains do not pay the cost they may pay under natural conditions when hosts may die before transmission.

Therefore, although serial passages are powerful tools to promote parasite adaptation, they poorly replicate the transmission conditions encountered in the wild. Our experimental design while being still far away from the natural conditions, at least allowed the cost of virulence to be expressed. Indeed, we found that parasitemia at the day of transmission increased over the passages and the lines that were finally lost (due to the host end point) were those that had the highest parasitemia levels. In agreement with this idea, we also found that, in the single infection treatment, once the most virulent lines were wiped out after ten passages, Py virulence decreased. Overall, these results nicely illustrate how host mortality prior to transmission can compromise any competitive advantage of fast replicating strains, which is one of the main assumptions of the trade-off model of parasite virulence (Cressler et al. 2016). Extrapolating results obtained under artificial lab conditions to the real world is never easy; however, with all the necessary caution, our results suggest that although conditions encountered in helminth coinfected hosts might favor the most virulent *Plasmodium* strains, reduced inter-host transmission due to mortality might rapidly offset the selective benefit of the virulent strains.

There are other alternative explanations for the lack of differences between SI-lines and COI-lines during the evaluation trials. First, it is possible that we did not passage Py long enough to observe a divergence between the lines. We stopped the passages for the COI-lines after seven passages because of the severity of the symptoms which resulted in the loss of 60% of the lines. That said, Py rapidly responded to the passages since the evolved lines (independently of the selection regime) had faster replication rate and induced more severe symptoms. Therefore, it is unlikely that the lack of divergence between the SI-lines and the COI-lines is due to the reduced number of passages. Another possible explanation is merely that Py encountered similar conditions in single infected and coinfected hosts, reducing the scope for selection to differentially operate on replication rate or virulence. However, previous work conducted on this (and similar) system(s) has consistently found that coinfected hosts have different immune profiles compared to single infections, both in terms of specific anti-malarial effectors and immunoregulatory pathways (Su et al. 2005, Hartgers and Yazdanbakhsh 2006, Noland et al. 2008, Tetsutani et al. 2009, Craig and Scott 2017, Kotepui et al. 2023, Dusuel et al. 2025). Therefore, it seems unlikely that Py experienced similar immune selection in single infected and coinfected hosts. That said, we cannot exclude that the strength of selection was not enough to produce a microevolutionary divergence between lines after seven passages.

Consistent with the results on the infection dynamics and virulence, we found that the infection with the evolved lines was equally tolerated by the host. We screened a couple of possible mechanisms accounting for variability in tolerance to malaria infection, the capacity to restore lost RBCs and the capacity to detoxify free heme (Chang et al. 2004, Kaiser et al. 2006, Seixas et al. 2009, Dey et al. 2014), and found that hosts infected with the COI-lines had a weaker HO-1 production compared to hosts infected with SI-lines. Although this result might suggest a weaker tolerance of hosts infected with COI-lines, the slope of the regression of RBC count on parasitemia was similar for hosts infected with SI-lines and COI-lines, showing that for a given intensity of the infection, the cost in terms of RBC loss was the same whatever the Py line. Interestingly, however, we found that evolved Py lines were less well tolerated by the host compared to the ancestral population. This result shows that the passages promoted Py genotypes that inflicted more damage to the host, independently from their capacity to replicate within the host. The mechanism underlying the reduced tolerance to the infection with the evolved lines deserves further investigation.

There have been few attempts to explore the microevolutionary response of a pathogen facing the conditions provided by a coinfected host. Leggett et al. (2013) let a bacteriophage to evolve under a single infection regime or a regime of mixed genotype infections and then assayed the evolved viral lines under the two environments (single or mixed genotype infections). The results showed that viral lines evolved in coinfected bacteria (mixed genotype infections) had a shorter time to lysis (i.e., killed the bacterium faster) compared to the single infection only when assayed under the mixed infection environment. This result is consistent with the idea that when competing with other viral strains, selection promoted time to lysis at the expense of viral yield (Legget et al. 2013). Although there are a number of major differences between this work and ours (coinfection involving different genotypes of the same species vs different species, type of interaction between competitors), we believe that the general take home message is that pathogens facing variable environmental conditions in single infected or coinfected hosts can rapidly adjust their strategy of host exploitation as to maximize their fitness.

In line with this argument, an important parameter shaping the fitness consequences following a selection episode is whether organisms face matched or mismatched environmental conditions. For instance, when pathogens are passaged in a novel host, they usually become adapted to this novel host (the matched environment) at the expense of the ancestral host (the mismatched environment) (Urbanowicz et al. 2016). We found evidence supporting this idea since Py lines passaged in single infected or coinfected hosts reached higher parasitemia when assayed in either single infected or coinfected hosts, compared to the COI-lines assayed in single infected hosts. While in agreement with the theoretical predictions, this result raises the question of the possible epidemiological consequences of the adaptation of *Plasmodium* parasites to helminth-coinfected hosts. In the intertropical zone, there is a large overlap between the areas of endemicity of *Plasmodium* and different species of gastrointestinal nematodes, resulting in high prevalence of coinfection, especially in the most vulnerable populations (Kepha et al. 2015, Degarege et al. 2016). *Plasmodium* therefore is likely to consistently experience both the environmental conditions provided by single infected and helminth-coinfected hosts, although other parameters, such as the order of infection, might change between rounds of transmission (Karvonen et al. 2019, Dusuel et al. 2025). When selection fluctuates over time (single infection, coinfection, etc.), plastic responses (as opposed to fixed strategies) are thought to confer the best fitness prospects, possibly maintaining exploitation rules allowing *Plasmodium* to maximize within-host replication and inter-host transmission according to the current environment provided by the host. Deworming campaigns are regularly deployed to protect exposed populations from the debilitating effects of helminths (Kepha et al. 2016, Wammes et al. 2016, Dila et al. 2022). However, deworming does not eradicate helminths from the entire population and therefore probably contributes to create fluctuating selection on coinfecting microparasites, possibly favoring the plastic adjustments of their host exploitation strategies.

## Acknowledgments

We are grateful to Valérie Saint Giorgio and all the staff of the animal facility for taking care of the animals.

## Author contribution

GS, BF, MR and BR conceived the study and gathered the funding; AD, LB, GS and BF collected the data; AD, LB, EG performed the lab work; GS analyzed the data; GS, AD and LB wrote the first draft of the manuscript; all authors revised and approved the final version.

## Data availability statement

All the data and codes are available in Zenodo (link).

